## Supplementary files for "The origins of placental mesenchymal stromal cells: Full spectrum flow cytometry reveals mesenchymal heterogeneity in first trimester placentae, and phenotypic convergence in culture"

### Titration of flow cytometry antibodies

All antibodies used in this work were titrated prior to use (Supplementary Figure 1 and 2). Titrations for Panel One are displayed in Supplementary Figure 1. Panel Two, designed for sorting villous core populations on a BD FACS Aria (a conventional cytometer), was optimized to spectrally spread antibodies out thereby reducing potential fluorophore spillover. The additional antibodies (conjugated to different fluorophores than those used in Panel One) were also titrated and are presented in Supplementary Figure 2. For some antibodies the optimal dose was difficult to detect due to the different autofluorescence and size of cells. Therefore, forward scatter was often employed to look at different sized cells, or CD45/β4 integrin were used to exclude hematopoietic and cytotrophoblast cells that interfered with detection of optimal antibody concentration in cell populations of interest (Supplementary Figure 3).

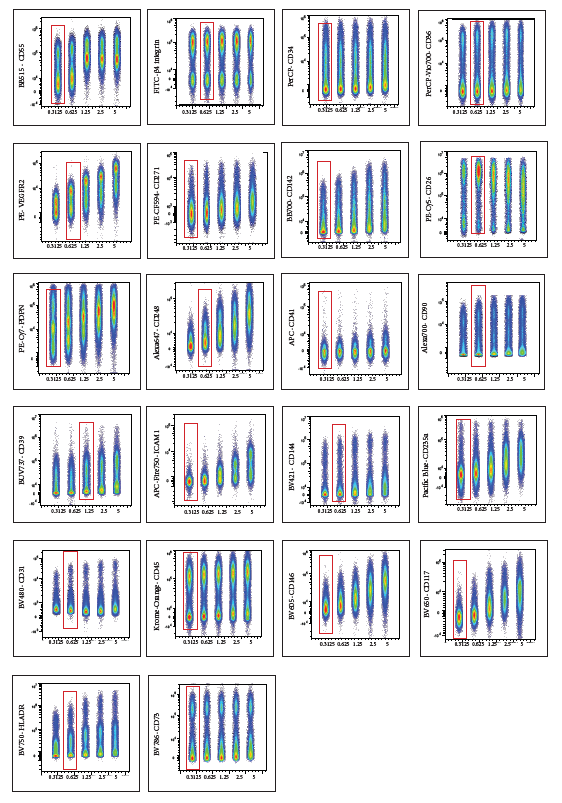

Supplementary Figure 1: All antibodies used in Panel One were titrated on placental villous core digest cells.

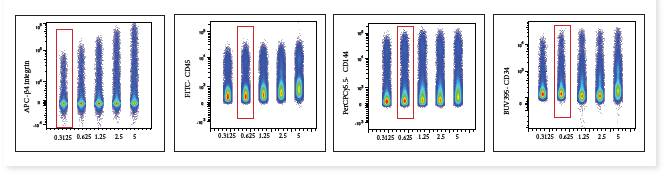

Supplementary Figure 2:Titration of additional antibodies, not contained in Panel One, required for the FACS sorting with Panel Two.

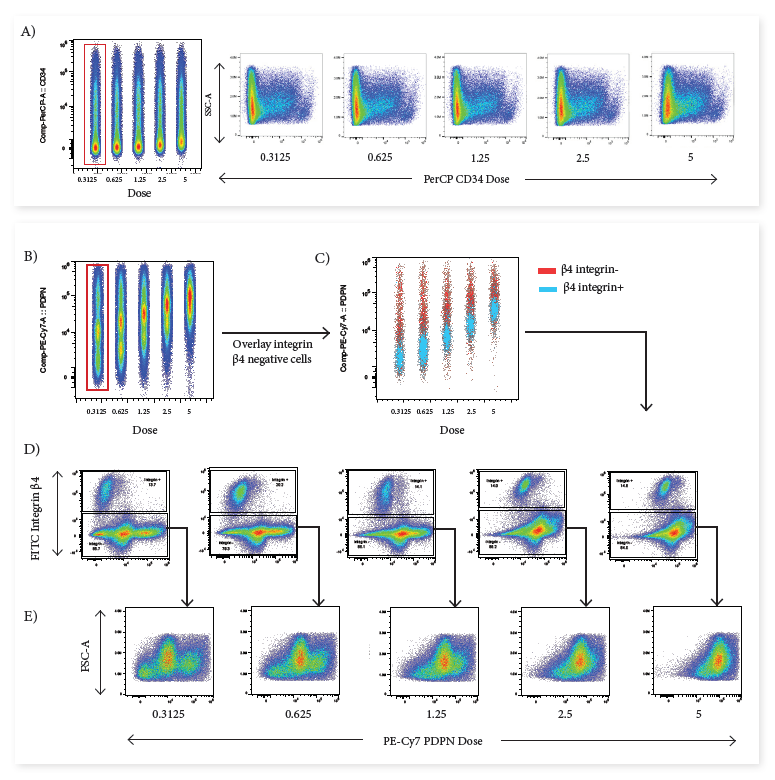

Supplementary Figure 3: Representative images depicting how forward scatter (FSC) (A) or addition of cell-specific antibodies improved detection of appropriate doses for specific placental populations. B) β4 integrin^+^ cells were negative for podoplanin but demonstrated an unspecific shift in expression at higher doses. C) Removal of β4 integrin^+^ improved detection of the optimal podoplanin dose.

### Analysis of CD73^+^CD90^+^ and podoplanin^+^CD36^+^ cells after 7 days in culture

After 7 days culture in EGM-2 medium on plastic tissue culture plates both CD73^+^CD90^+^ and podoplanin+CD36+ cells expressed the basic “MSC” phenotype (CD73^+^CD90^+^CD31^-^CD144^-^CD45^-^HLADR^-^) (Supplementary Figure 4). Podoplanin+CD36+ remained very homogeneous for the markers assessed (podoplanin^+^CD142^+^CD26^+^) however, they downregulated expression of CD36 while upregulating expression of CD146. Conversely CD73+CD90+ cells demonstrated phenotypic (podoplanin, CD26 and CD146) differences between samples and heterogeneity within samples (Supplementary Figure 4). Explant cultured pMSCs were assessed in order to determine how their phenotype compared with CD73^+^CD90^+^ cells (Supplementary Figure 5). All pMSCs were CD73^+^CD26^+^podoplanin^+^CD142^+^CD146^+^ aligning more closely with the podoplanin^+^CD36^+^ cells. As previously demonstrated pMSCs cultured in EGM-2 downregulated CD90. Summary data for these results are displayed in the main manuscript (Figure 4), but here for transparency histograms of individual samples are provided.

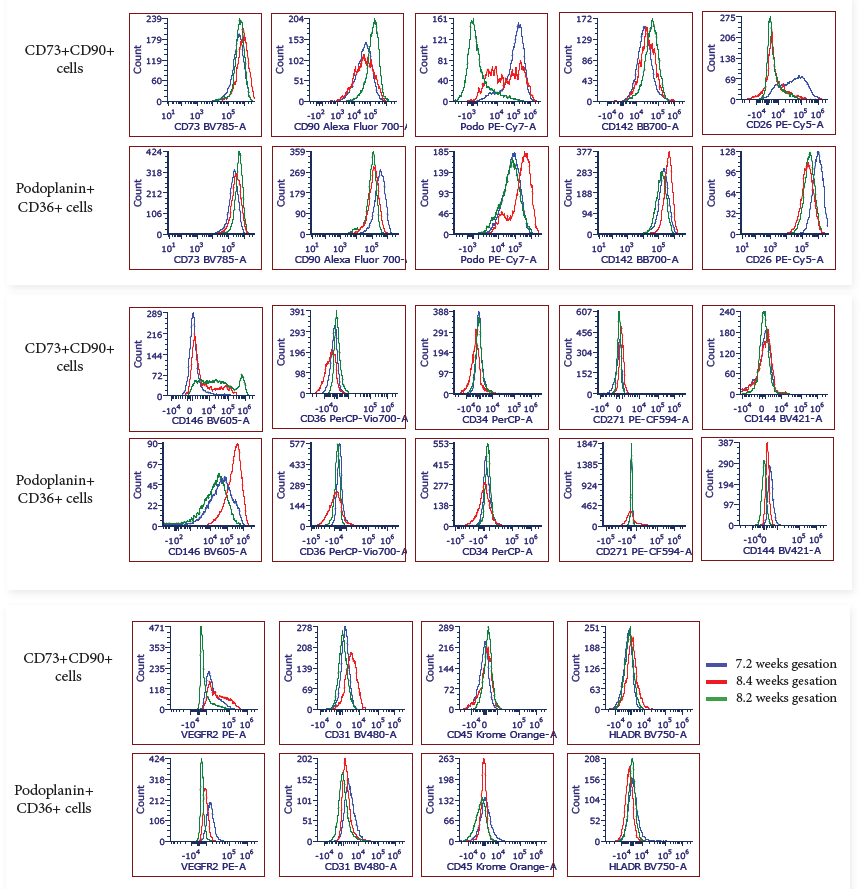

Supplementary Figure 4: Flow cytometry histograms displaying the phenotype of FACS sorted CD73^+^CD90^+^ and podoplanin^+^CD36^+^ cells after 7 days culture *in vitro* (n=3).

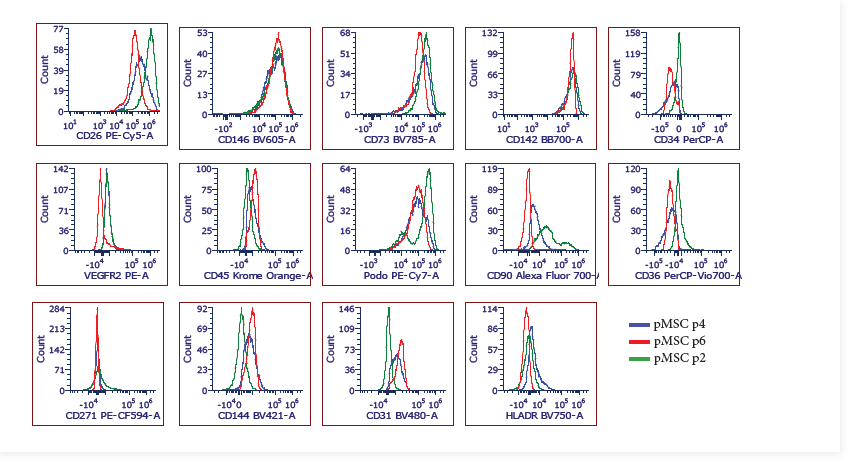

Supplementary Figure 5: Flow cytometry histograms displaying the phenotype of explant isolated pMSCs after culture *in vitro* (n=3, p2-6)..
